# Spatially clustered pattern of transcription factor binding reveals phase-separated transcriptional condensates at super-enhancers

**DOI:** 10.1101/2023.06.18.545510

**Authors:** Zhenjia Wang, Shengyuan Wang, Chongzhi Zang

## Abstract

Many transcription factors (TFs) have been shown to bind at super-enhancers, forming transcriptional condensates to activate transcription in many cellular systems. Genomic and epigenomic determinants of phase-separated transcriptional condensates are not well understood. Here we systematically analyzed DNA sequence motifs and TF binding profiles across human cell types to identify the molecular features that contribute to the formation of transcriptional condensates. We found that most DNA sequence motifs are not distributed randomly in the genome, but exhibiting spatially clustered patterns associated with super-enhancers. TF binding sites are further clustered and enriched at cell-type-specific super-enhancers. TFs exhibiting clustered binding patterns also have high liquid-liquid phase separation abilities. Compared to regular TF binding, densely clustered TF binding sites are more enriched at cell-type-specific super-enhancers with higher chromatin accessibility, higher chromatin interaction, and higher association with cancer outcome. Our results indicate that the clustered pattern of genomic binding and the phase separation properties of TFs collectively contribute to the formation of transcriptional condensates.

## INTRODUCTION

Transcription factors (TFs) play essential roles in driving transcriptional activation by binding at DNA and inducing cell type-specific promoter-enhancer interactions in the genome^1, 2^. TF activities are important in numerous biological processes and transcriptional dysregulation has been found to associate with many diseases such as cancer^3^. Super-enhancers (SEs) are a special type of enhancer-like ultra-broad genomic regions which exhibit strong and broad enrichment of mediator and enhancer-associated histone marks such as H3K27ac^4–6^. An SE usually contains multiple cis-regulatory (enhancer) elements and is bound by multiple TFs. The enhancer sequences, which contain the short DNA motifs recognized by DNA-binding TFs, act as platforms to recruit gene control machinery including the TFs and co-activators at specific genomic loci^7^. SEs as clusters of enhancers that are occupied by high-density of TFs can drive higher levels of transcription than typical enhancers^5^. Active SEs have been observed in cancer cells^6, 8^, stem cells^4, 9^, and normal somatic cells^5, 10^.

Liquid-liquid phase separation (LLPS) and the formation of transcriptional condensates are implicated as potential mechanisms of SEs^11–13^. The activation of functional enhancers/SEs requires the binding of both cell-type specific factors and sequence-dependent effectors to drive the formation of localized condensation and promote enhancer activity and transcription^14, 15^. Multiple TFs including CCCTC-binding factor (CTCF) may involve in this process with either driving or instrumental functions^16^. TFs, mediator, and RNA polymerases II have been found to form clusters in the cell nucleus^17, 18^, indicating the formation of phase-separated condensates. LLPS and condensate formation usually require a large aggregation of protein molecules with intrinsic disordered domains (IDRs)^19^. The LLPS ability of a protein can be quantitatively characterized by its sensitivity to 1,6-hexanediol (1,6-HD) treatment, which can disrupt the LLPS condensates in vitro and in vivo^20^. An anti-1,6-HD index of chromatin-associated proteins (AICAP)^20^ has been used to quantify the LLPS ability of thousands of nuclear proteins^20^. Proteins with low AICAP (between 0 and 1) are associated with high content of IDRs and high LLPS potential.

TF binding patterns are determined by both DNA sequence^21^ and cell type-specific chromatin structure and accessibility. TFs can function to regulate target genes at various spatial ranges in the genome^22^. The spatial distribution of TF binding sites across the genome has been briefly examined using ChIP-chip data but not extensively surveyed with the more recently available high-throughput sequencing data^23^. TF hotspots have been observed where many TFs colocalize in narrow regions in the genome^24, 25^. However, to what extent the genomic distribution of TF motif-matching DNA sequences and TF binding sites affect the activities of SEs and the formation of transcriptional condensates globally, and what genomic features can influence condensate formation at specific genomic loci, are poorly understood. Most existing nuclear LLPS/condensate studies did not use the rich genomic data, while genomics studies on SEs are difficult to connect to LLPS/condensate phenomena. There is a pronounced gap between data-driven predictions from genomics perspective and the experimental studies of transcriptional condensate formation.

In this study, we performed a comprehensive survey of 528 human TFs’ known sequence motifs and 6,650 ChIP-seq datasets in a variety of human cell types, and developed a statistical metric to quantify the genomic clustering pattern of TF binding. We found that most TFs’ motif matching sites and in vivo binding sites both exhibit a spatially clustered pattern in the genome. Clustered motif sites and clustered TF binding sites are enriched at super-enhancers. We found that the clustering tendency of TF binding is correlated with TF’s LLPS property measured by AICAP. By integrating the TF binding profiles in colorectal cancer and breast cancer with molecular genomic profiling data from The Cancer Genome Atlas (TCGA), we identified cancer-specific clustered TF binding sites and found a significant association with cancer patient survival, indicating the functional importance of transcriptional condensates in cancer.

## RESULTS

### Clustered TF motif sites are enriched at putative super-enhancers

To get a comprehensive survey of spatial distribution patterns of cis-regulatory elements in the genome that are potential TF binding sites, we collected 528 human TF sequence motifs from the Jaspar database^26^ and 6,650 high-quality ChIP-seq TF binding profiles from the Cistrome database^27^. For each TF motif, we used FIMO^28^ to identify its genome-wide motif matching sites (TFMSs) and examine their location distribution in the genome (Fig. 1a). To quantify the spatial clustering tendency of the genomic distribution pattern of a TFMS, we generated a control by placing the same number of genomic loci randomly in the genome, following the Poisson point process. We define a metric, cluster propensity (CP), as the two-sided Kolmogorov-Smirnov (K-S) test statistic between the genomic interval distribution of the TFMSs and that of the control, to quantify the genomic clustering tendency of a TFMS profile (Fig. 1a). Intuitively, a TFMS profile with a spatially clustered pattern will have a positive CP (Fig. 1b,c). If the TFMS interval distribution is modeled by the Gamma distribution^23^, the CP is correlated with the shape parameter *k* in the Gamma distribution (Supplementary Fig. 1a). TFMS CP is not correlated with the total number of motif matching sites in the genome, or the motif sequence length (Supplementary Fig. 1b-d), indicating the robustness of this metric. Among the 528 TFs analyzed, 417 (79%) show a positive CP, indicating the TFMSs are more clustered than random in the genome (Fig. 1d). The motif matching sites of the TFs with high TFMS CP are significantly enriched at the union of super enhancers (SEs) (Fig. 1d, with examples at Fig. 1e, p < 0.05, by Fisher’s exact test). CENPB, a centromere protein, has the highest TFMS CP across all TFs (Fig. 1b), and EWSR1-FLI1, which recruits BAF complexes to tumor-specific enhancers and activates transcriptional events of Ewing’s sarcoma^29^, also ranks on top with high TFMS CP (Fig. 1c). These results suggest that most TFs’ sequence motif matching sites have a higher clustering tendency than randomly distributed in the genome.

**Figure 1.**
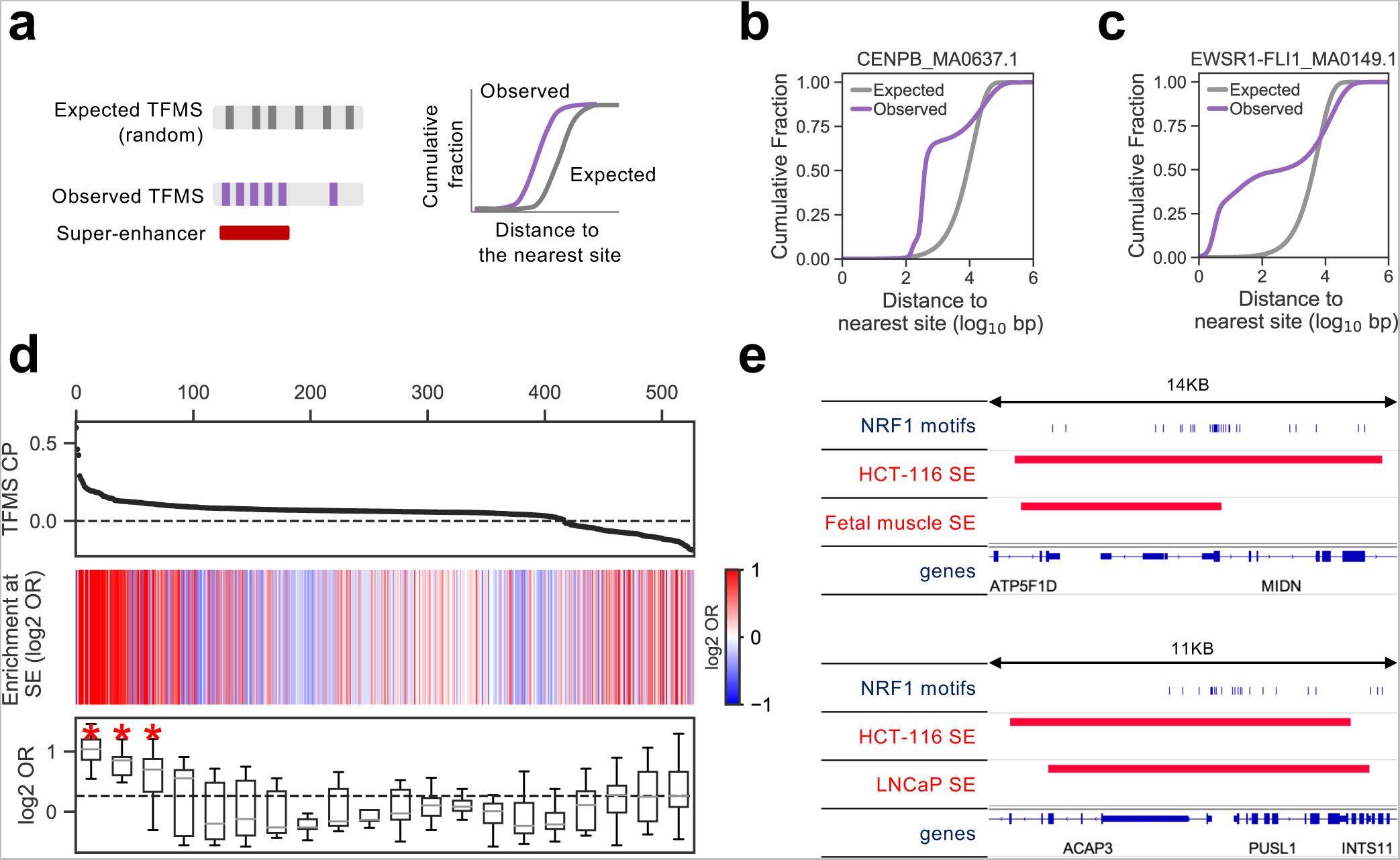
Clustered transcription factor motif sites (TFMS) are enriched at super-enhancers (SEs). **(a)** Schematic of TFMS cluster propensity (CP). K-S test is used to compare the cumulative distributions of distance to the the nearest downstream site between the TFMS profile (Observed) and the random control (Expected). **(b,c)** Cumulative distributions of distance to the nearest downstream motif site for CENPB (b) and EWSR1-FLI1 (c) and their corresponding control (expected random distribution). **(d)** Association of TFMS CP with their enrichment at union SEs. Top: Rank of 528 TF motifs by TFMS CP. Middle: Enrichment (log2 odds ratio) of each TFMS profile at union SEs compared to genomic control. Bottom: The 528 motifs were divided into 20 equal-size groups. The boxplots show the enrichment (log2 odds ratio) of TFMS at union SE compared to genomic control. * p<0.05, by one-sample one-sided t-test. **(e)** Genome browser snapshots of NRF1 motifs and the surrounding SEs.

### Clustered TF binding sites are enriched at cell type-specific super-enhancers

DNA sequence only provides the basic anchors of potential TF binding but is not sufficient to determine the actual binding profile of a TF in a cell type. Therefore, we next examined the 6,650 high-quality ChIP-seq binding profiles to evaluate the clustering tendency of actual TF binding sites (TFBSs). With the assumption that most TFBSs contain a motif matching sequence, for a TF binding profile containing a number of binding sites, we randomly sampled the same number of motif sites from the TFMS profile as the control (Fig. 2a). Similarly, we defined the TFBS CP as the two-sided K-S test statistic between the genomic interval distribution of the TFBSs and that of the control, to quantify the genomic clustering tendency of a TFBS profile (Fig. 2a). The TFBS CP is also a robust metric that is not sensitive to the number of binding sites called from ChIP-seq data (Supplementary Fig. 2). Interestingly, we found that all the top 20 TFs mostly shared across 6 cell types exhibit a positive TFBS CP, indicating a high clustering tendency (Fig. 2b), and these TFBSs are enriched at cell-type specific SEs compared to genomic control (Fig. 2c). Furthermore, the TFBS CP of a TF profile is highly correlated with the TF profile’s enrichment level at SEs, demonstrating a strong association between the spatially clustered TF binding pattern and SEs (Fig. 2d). Considering TFBSs may occur at genomic regions without sequence motifs, we checked the CP of TFBS with or without sequence motifs and found that the TFBSs without motifs even have a higher CP and higher enrichment at cell-type-specific SEs compared to TFBSs with motifs (Supplementary Fig. 3a-c). We found different TFs show different TFBS CP and different enrichment levels at SEs within the same cell type (Supplementary Fig. 3d), while the same factor also shows different TFBS CPs and different enrichment levels at SEs across different cell types (Supplementary Fig. 4), indicating the cell-type specificity of TF binding.

**Figure 2.**
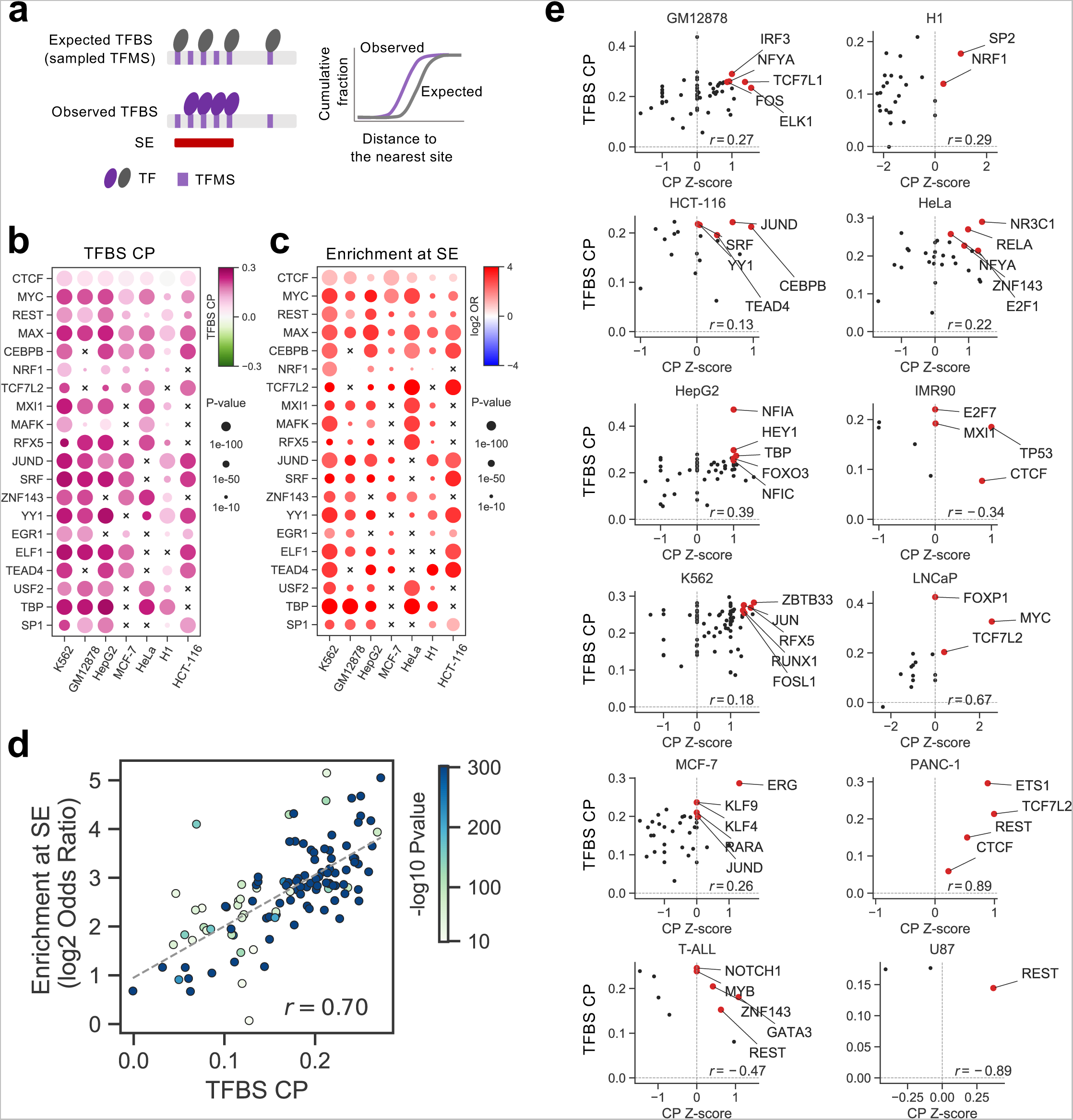
Clustered transcription factor binding sites (TFBS) are enriched at cell type-specific super-enhancers (SEs). **(a)** Schematic of TFBS CP. K-S test is used to compare the cumulative distributions of distance to the the nearest downstream site between a TFBS profile (Observed) and the random control (Expected), generated by randomly selecting the same number of motif sites. **(b)** TFBS CP of 20 TFs in 6 cell types. The color indicates TFBS CP and the circle size indicates p-value calculated by K-S test. **(c)** Enrichment of TFBS at cell type-specific SE compared with random control (expected). The color indicates the enrichment at SE (log2 odds ratio) and the circle size indicates p-value calculated by the Fisher’s exact test. **(d)** Scatter plots of profiles for 20 TFs in 6 cell types for TFBS CP (x-axis) and their enrichment at cell type-specific SEs compared with random control (y-axis). **(e)** Scatter plots of TFs showing their TFBS CP (y-axis) and z-scaled TFBS CP (x-axis) in each of the 12 cell types with at least 3 TFs having ChIP-seq data.

**Figure 3.**
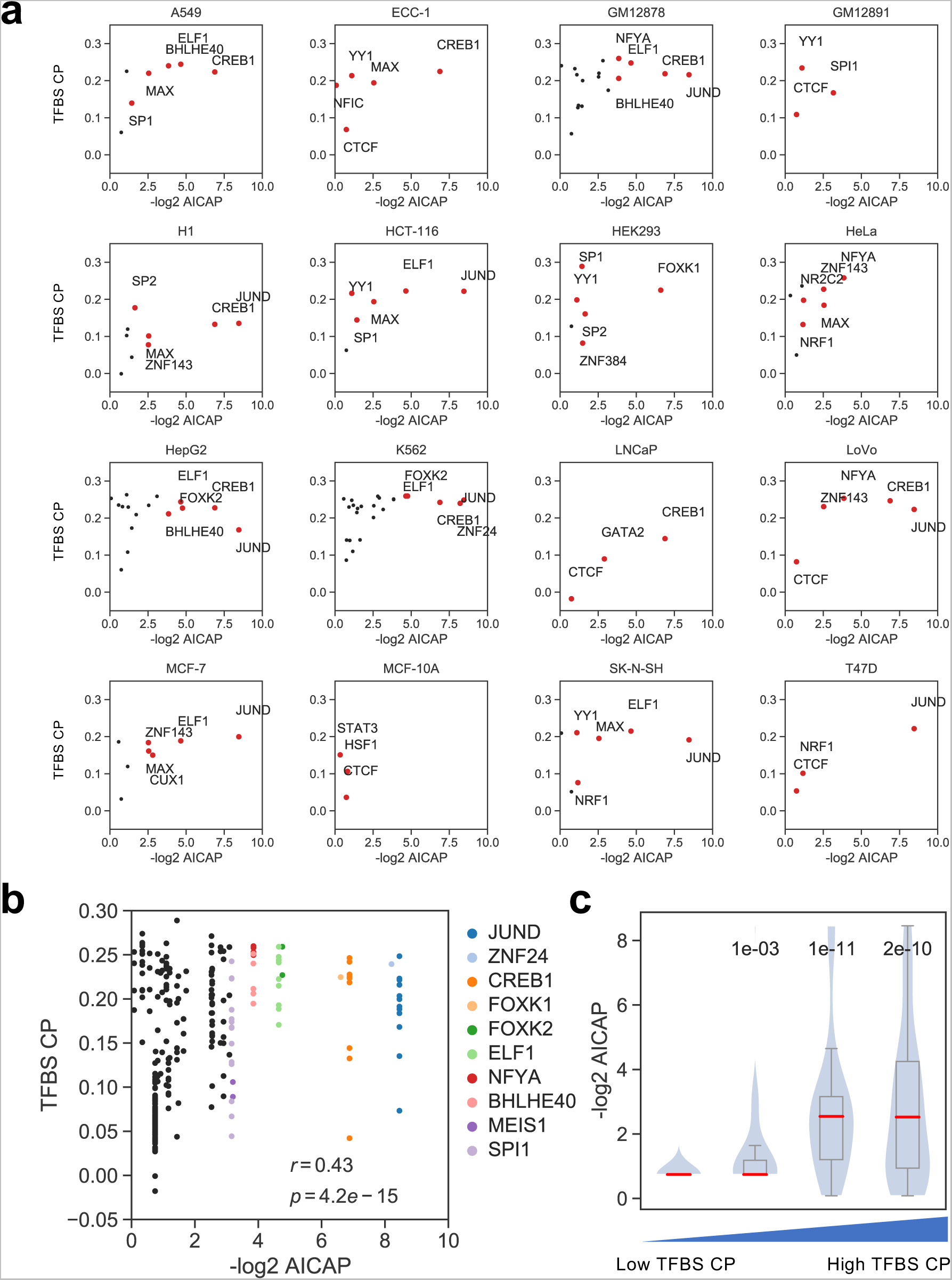
Clustered transcription factors are associated with LLPS potential. **(a)** Scatter plots of TFBS CP (y-axis) against -log2 AICAP score (x-axis) in 9 cell types, each of which has at least 3 TFs with both ChIP-seq and AICAP data available. A lower AICAP score (higher –log2 AICAP) indicates a higher potential of liquid-liquid phase separation (LLPS). **(b)** Scatter plots of TFBS CP (y-axis) against log2 AICAP score (x-axis) of all TFs across all cell types with both ChIP-seq and AICAP data available. **(c)** Box plots of -log2 AICAP scores for 4 quartiles of TFs grouped by TFBS CP. Numbers in the plot are the p-values comparing the -log2 AICAP scores in the corresponding quartile with the first quartile, calculated by the one-sided Student’s t-test.

**Figure 4.**
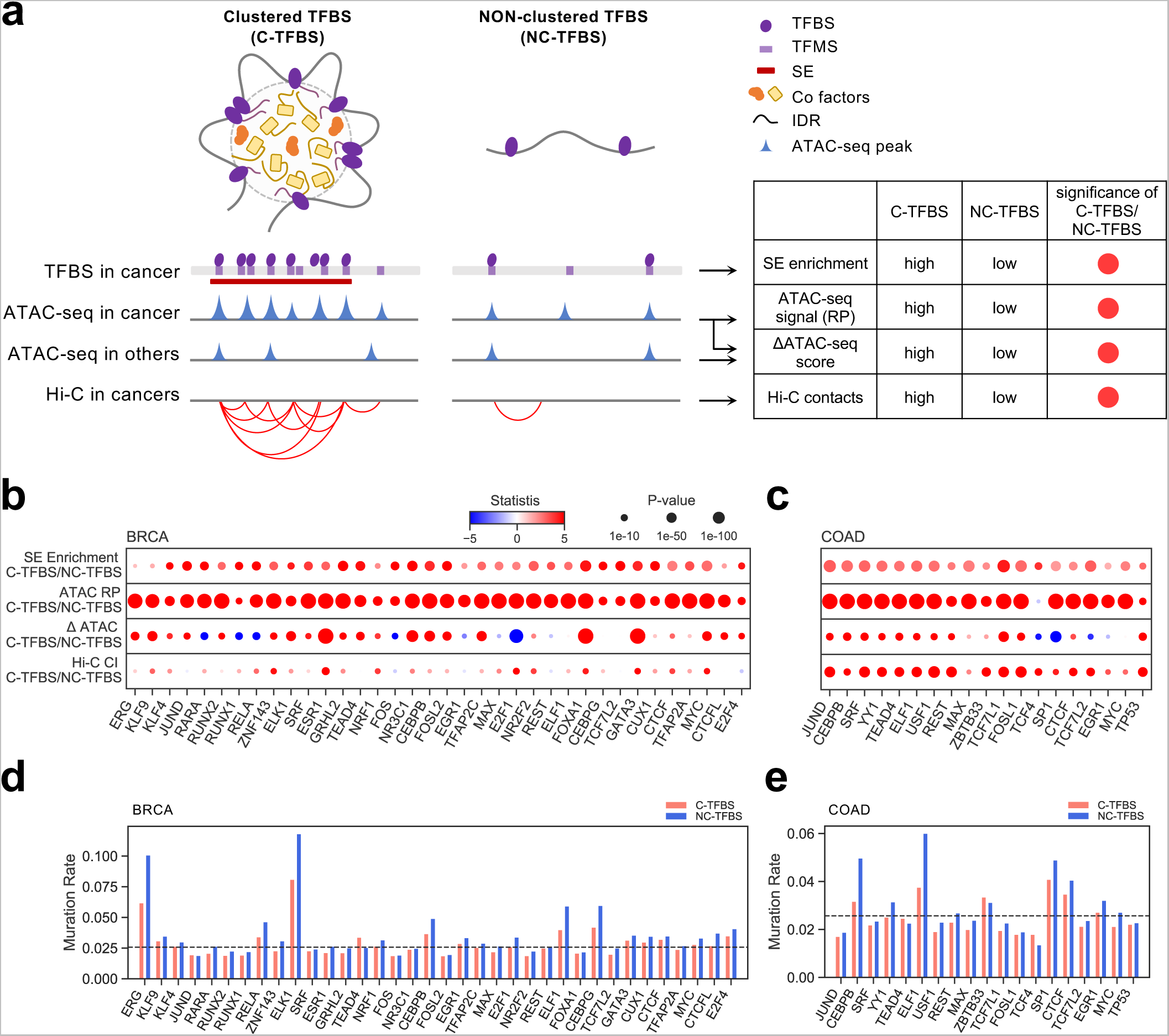
Clustered TFBS show higher SE enrichment and higher chromatin activities in cancer cells. **(a)** Schematic of the epigenomic features comparing between clustered (C-) and non-clustered (NC-) TFBS. **(b,c)** C-TFBS and NC-TFBS comparison in cell-type-specific SE enrichment, ATAC-seq RP, differential ATAC-seq score, and Hi-C interactions, in BRCA (b) and COAD (c). TFs were ranked along the x-axis by CP rank (average rank of TFBS CP and z-scaled CP) as shown in Fig. 2e. **(d,e)** Mutation rate at motif loci within the binding sites comparing C-TFBS and NC-TFBS in BRCA (d) and COAD (e). TFs were ranked along the x-axis by CP rank (average rank of TFBS CP and z-scaled CP) as shown in Fig. 2e.

We next used both the absolute and the normalized TFBS CPs to identify potential key factors with high cell-type specific CPs in each cell type (Fig. 2e). We identified JUND on the top of the list for several cell types including the colon cancer cell line HCT-116 and the breast cancer cell line MCF7, while JUND overexpression increases the cell proliferation in prostate cancer^30^ and enhanced JunD signaling is responsible for BET inhibition resistance in cancers^31^. NFIA was shown as the top ranked TF in the liver cancer cell line HepG2 and was indeed overexpressed in various cell lines including HepG2^32^. MYC, the top ranked TF in the prostate cancer cell line LNCaP, is overexpressed and associated with poor survival in human prostate cancer and has been shown as a major driver of prostate cancer tumorigenesis^33, 34^. ERG, the top ranked TF in the breast cancer cell line MCF7, can induce a mesenchymal-like signature and is positively correlated with invasive breast cancer^35, 36^. ETS-1 is the top ranked factor in the pancreatic cancer cell line PANC-1 and is overexpressed in pancreas^37^ while its increased binding activity is critical for PANC-1 cellular invasiveness^38^. NOTCH1 and GATA3 were shown on top in T-ALL. NOTCH1 is a major oncogenic TF in T-ALL^16, 39^, and GATA3-mediated enhancer nucleosome eviction was shown as a driver of MYC expression and is strictly required for NOTCH1-induced T-ALL initiation and maintenance^40^. These results suggest that many TF binding sites show a further clustering tendency on top of motif sites with an enrichment at cell-type-specific SEs, and that a TF’s high cell type specific CP can be indicative of its important oncogenic functions in cancer cells.

### Transcription factors with highly clustered binding have high liquid-liquid phase separation potential

The association between clustered TF binding and SEs reminded us of the possible phenomena of transcriptional condensate formation contributed by TF proteins. To determine other potential factors that contribute to the clustered pattern of TF binding in addition to DNA sequences, we next examined the liquid-liquid phase separation (LLPS) property of TF proteins. In 16 cell types with most TF ChIP-seq profiles^20^, we found a subtle but clear trend that the TFs with higher TFBS CP tend to have lower AICAP (Fig. 3a), indicating their higher ability to form phase separated condensates in cells. Remarkably, putting together 300 binding profiles of 30 different TFs in 154 cell types, we found a significant correlation between TFBS CP and AICAP (Fig. 3b). If we grouped all TFBSs into four quartiles based on their TFBS CP, we could see that the negative log-transformed AICAP of the TFs in the third and fourth quartiles with the highest TFBS CPs are significantly higher than that in the first and second quartiles (Fig. 3c). These results indicate that the intrinsic LLPS property of TF protein molecules might contribute to the formation of phase-separated transcriptional condensates at SEs. LLPS of TF proteins that contain intrinsically disordered regions (IDRs) might be a driver of transcriptional condensate and super-enhancer formation.

### Clustered TFBSs show active chromatin features and higher enrichment at SEs in cancer cells compared to non-clustered TFBSs

Besides using the CP metric to quantify the global feature of a TF binding profile, we also characterized the genomic regions with densely clustered binding sites of a TF and compared with those binding sites that are not clustered in the genome in cancer. We defined the clustered TFBSs (C-TFBSs) as those that are significantly closer to its nearest binding site than expected in the control distribution, and called the remaining sites non-clustered TFBSs (NC-TFBSs) (Fig. 4a). Integrating the genome-wide chromatin accessibility profiling (ATAC-seq) data from The Cancer Genome Atlas (TCGA)^41^ with publicly available data such as 3D genome Hi-C maps and SE annotations from matched cancer types, we compared the chromatin accessibility, chromatin interaction and cell-type-specific SE enrichment between C-TFBSs and NC-TFBSs in breast cancer (BRCA), colon cancer (COAD), cervical cancer (CESC), liver cancer (LIHC), and prostate cancer (PRAD), where data for the matched cancer cell types exist.

We found that all TFs’ C-TFBSs are significantly enriched at cell-type specific SEs compared to NC-TFBSs for all cancer types examined (Fig. 4b,c, Supplementary Fig. 5a) (p < 0.05, by Fisher’s exact test). We quantified the ATAC-seq signal at each TFBS using the regulatory potential (RP) metric^42^ for comparison between C-TFBSs and NC-TFBSs, and found that the C-TFBSs show significantly higher (p < 0.05, by two-tailed Student’s t-test) RPs compared to NC-TFBSs for all TFs in all cancer cell types, indicating a higher chromatin accessibility level at C-TFBSs (Fig. 4b,c, Supplementary Fig. 5a). Meanwhile, we calculated the differential ATAC-seq signals in each cancer type comparing to other samples from all other cancer types as control and found that the C-TFBSs show significantly higher differential chromatin accessibility compared to NC-TFBSs for the vast majority of TFs (Fig. 4b,c, Supplementary Fig. 5a) (p < 0.05, by two-tailed Student’s t-test). We also found that the C-TFBSs tend to have significantly higher chromatin interactions with their surrounding genomic regions compared to NC-TFBSs (Fig. 4b,c, Supplementary Fig. 5a) (p < 0.05, by two-tailed Student’s t-test). These results indicate that those genomic regions with highly clustered TF binding are more active with higher chromatin accessibility, higher chromatin interactions and higher enrichment at SEs compared to genomic regions with NC-TFBSs.

**Figure 5.**
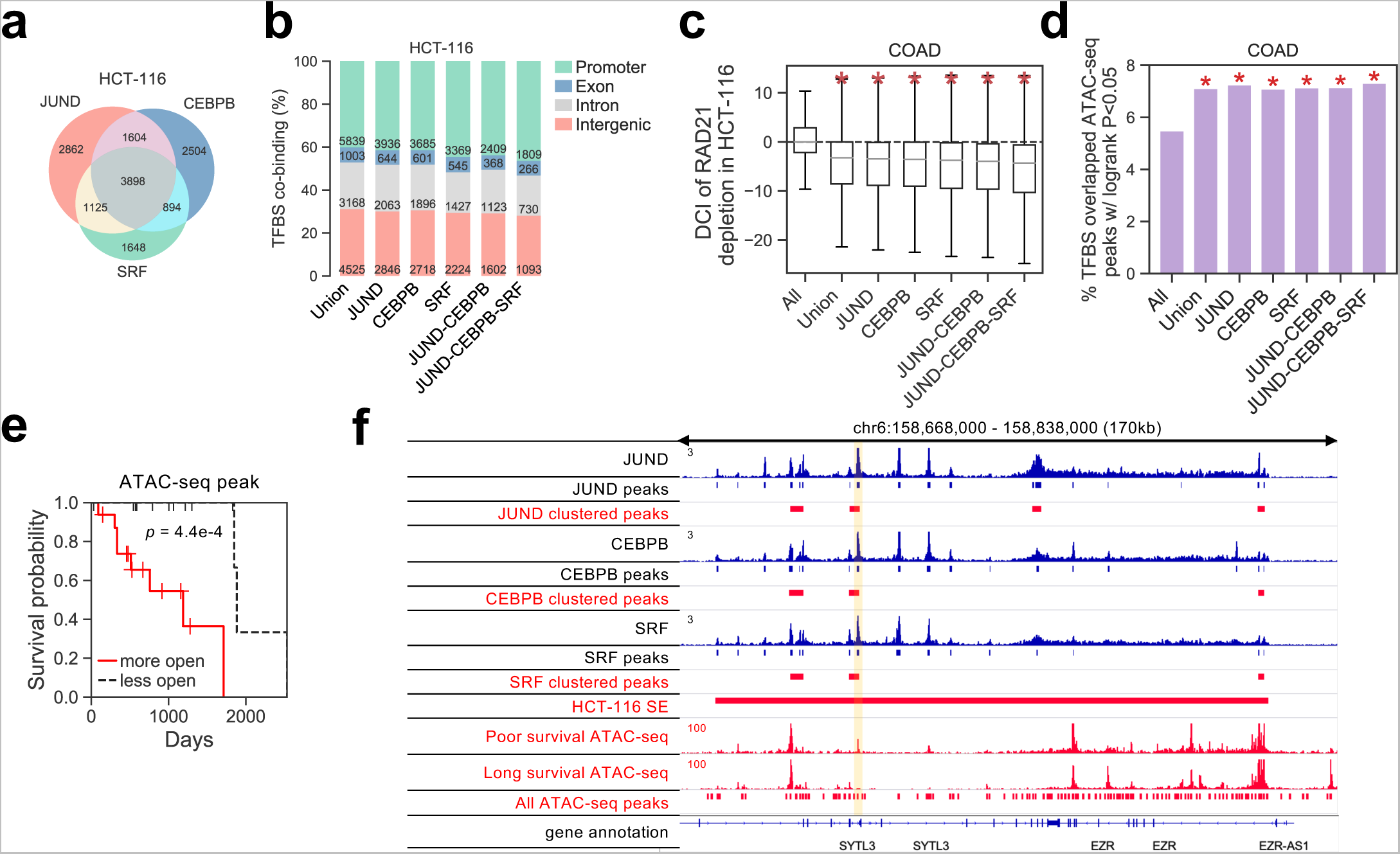
Chromatin accessibility at clustered TF co-binding sites is predictive of COAD survival. **(a)** Numbers of co-binding of clustered sites of JUND, CEBPB and SRF, the 3 factors with the highest ranked CP in COAD. **(b)** Genomic distributions of binding and co-binding of of the 3 factors’ clustered sites. **(c)** Differential chromatin interaction (DCI) levels at binding and co-binding of the 3 factors’ clustered sites. DCI were calculated comparing before and after RAD21 degradation in HCT-116 cells. * p<0.05, by two-sided Student’s t-test. **(d)** Percentage of ATAC-seq peaks overlapping with each category that are significantly associated with COAD survival. * p<0.05, by two-sided Student’s t-test. **(e)** Univariate survival analysis comparing patients with high (red) and low (black) chromatin accessibility at the clinical-associated ATAC-seq peaks. P-value by log-rank test. **(f)** Example ChIP-seq and ATAC-seq signals surrounding an ATAC-seq peak.

The DNA binding TFs are highly specific to the presence of its binding sequence motif and can be compromised by mutations affecting the consensus motif sequence^43^. We analyzed the whole-genome sequencing (WGS) data from BRCA, CRC, CESC, LIHC and PRAD patient samples from the International Cancer Genome Consortium (ICGC)^44^, but did not see significantly higher mutation rate at the sequence motif matching site within C-TFBSs compared to NC-TFBSs across all TFs in any cancer type (p > 0.05, by the two-tailed Student’s t-test), and very few TFs show a higher mutation rate in their binding motif sites than the average mutation rate in the genome (Fig. 4d,e, Supplementary Fig. 5b). We next examined whether the mutations of genes encoding the TFs potentially associate with transcriptional condensates at the TFBSs. We separated the patient samples in each cancer type into two groups by the ATAC-seq RPs at the C-TFBSs to mimic those samples that contain transcriptional condensates and others. However, we did not see any significant difference in TF gene mutations between the samples with high C-TFBS RP and others with lower RPs (Supplementary Fig. 6). These results suggest that the majority of cancer patient-specific clustered TFBSs are not due to DNA mutations altering the consensus binding sequence.

**Figure 6.**
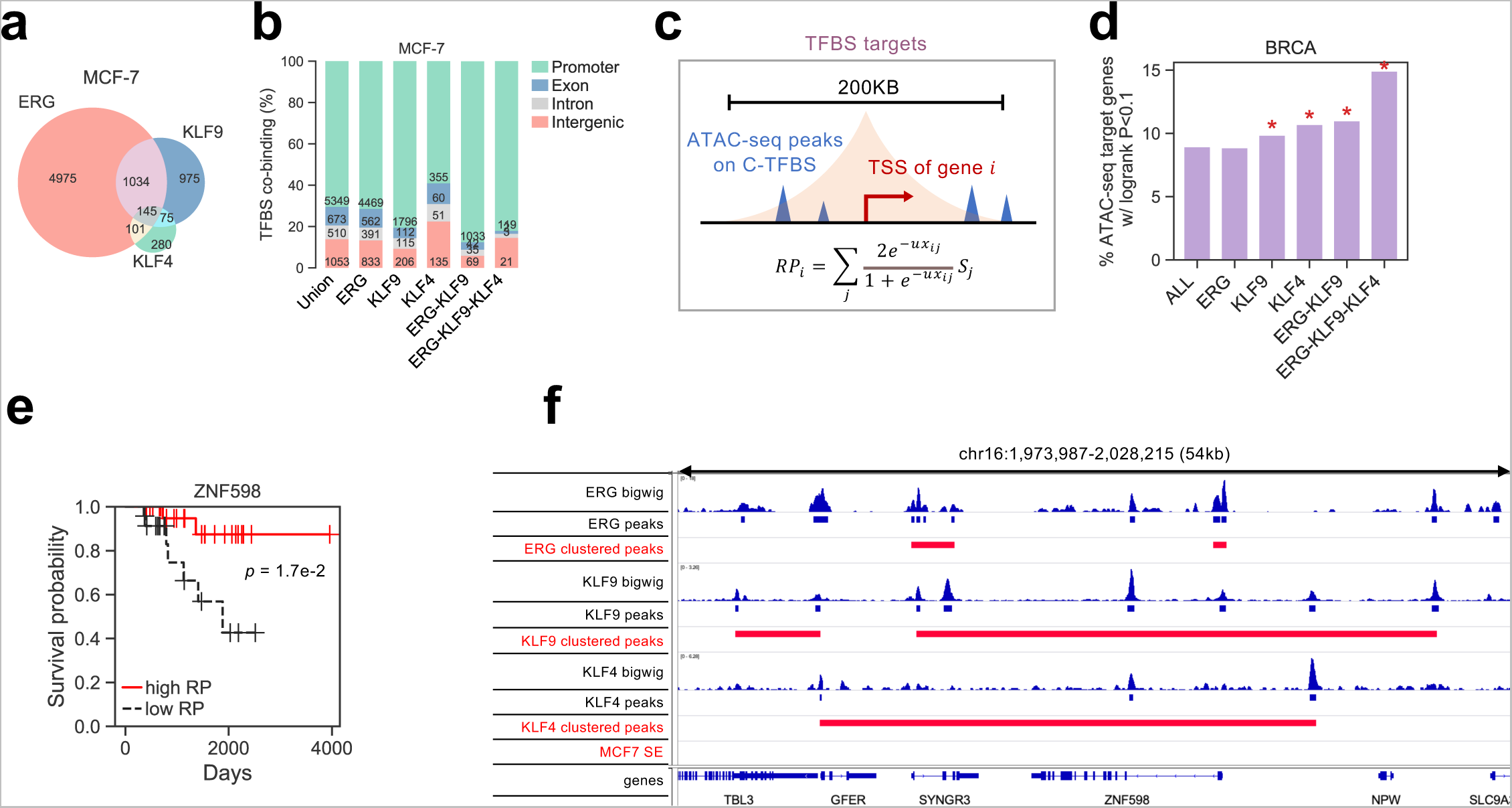
Co-regulated genes of clustered TFBSs are predictive of BRCA survival. **(a)** Venn diagram of co-binding of clustered sites of ERG, KLF9, and KLF4, the 3 factors with the highest ranked CP rank (average rank of TFBS CP and z-scored CP) in BRCA. **(b)** Genomic distributions of binding and co-binding of the 3 factors’ clustered sites. **(c)** Schematic of TF regulatory potential (RP) on target genes. Identified TFBSs overlapped ATAC-seq peaks surrounding a gene locus (TSS+/-100KB) were collected and the weighted sum was calculated as the RP for this gene. **(d)** Percentage of the target genes of each category that are significantly associated with BRCA survival. * p<0.05, by two-sided Student’s t-test. **(e)** Univariate survival analysis at gene ZNF598 comparing patients with high (red) and low (black) ATAC-seq RP. P-value was identified by log-rank test. **(f)** Example of ChIP-seq and ATAC-seq signals surrounding the gene ZNF598.

### Chromatin accessibility levels at clustered TF co-binding sites are predictive of COAD survival

Assuming the C-TFBSs have higher transcriptional activity with higher chromatin accessibility and chromatin interactions than NC-TFBSs, we then sought to study whether the C-TFBSs are functionally important in cancer cells and their potential relevance to clinical outcome. We focused on two cancer types, COAD and BRCA, considering they have sufficient samples with clinical data in TCGA. We used the top 3 TFs, JUND, CEBPB, and SRF, with the highest ranked TFBS CP in HCT-116 cells, to study the potential functions of C-TFBSs in COAD. Interestingly, among the total of 14,535 union C-TFBSs of the three factors, 3,898 (27%) are co-occupied by all three TFs (Fig. 5a), and over 19% and 28% of the co-binding sites are in the intronic or intergenic regions, respectively (Fig. 5b). We next used dynamic Hi-C data in HCT-116 cells before and after RAD21 degradation, in which promoter-enhancer interactions and chromatin condensates were disrupted, to characterize the differential chromatin interactions (DCI) in the genome^45^. We found that the C-TFBSs of JUND, CEBPB, and SRF and the co-binding regions exhibited significantly decreased chromatin interactions with their surrounding genomic regions (<100kb) after RAD21 degradation (Fig. 5c) (p < 0.05, by two-tailed Student’s t-test). Putting together, the high co-localization, high occurrence at non-coding regions, and high enrichment at SEs, suggest that the clustered co-binding regions of the three factors are likely associated with transcriptional condensates in colon cancer.

We next accessed how the co-binding regions of the C-TFBSs are associated with patient survival. We performed univariate survival analysis for each union chromatin accessibility region using ATAC-seq data from TCGA COAD samples. We found the ATAC-seq peaks overlapped with the clustered binding sites of JUND, CEBPB, and SRF and their co-binding regions are significantly more likely to be associated with survival than a random ATAC-seq peak from the genome (Fig. 5d) (p < 0.05, by Fisher’s exact test). At 66% of the co-binding regions a high chromatin accessibility level would significantly associate with poor survival (p < 0.05, by log-rank test), shown in Fig. 5e as an example. An example of survival-associated ATAC-seq peaks co-bound by the three TFs in a super-enhancer region is shown in Figure 5f.

### Co-regulated genes of clustered TFs are predictive of BRCA survival

Unlike COAD, the 3 TFs, ERG, KLF9, and KLF4, with the highest CP rank in breast cancer cell line MCF7 do not co-occupy their C-TFBSs significantly. Among the total of 7,585 union C-TFBSs, only 145 (1.9%) are co-occupied by all three factors (Fig. 6a), most (82%) of which are at gene promoters (TSS+/-2kb) (Fig. 6b). The survival analysis using the ATAC-seq data from the TCGA BRCA samples do not show significant association between the chromatin accessibility level at C-TFBS co-binding regions and patient survival (Supplementary Fig. 7a). Considering the enrichment of the C-TFBS co-binding regions at gene promoters, we sought to examine the putative target genes of the three factors. We calculated the RP score of the ATAC-seq peaks overlapped with a set of TFBSs or co-binding sites to each gene. The target genes of each TF or co-binding sites were selected as those with RP ≥ 0 (Fig. 6c). We performed univariate survival analysis for each gene using ATAC-seq RP, and found the target genes of KLF9, KLF4 and the co-targets are all significantly associated with survival (Fig. 6d). For example, the three factors ERG, KLF9 and KLF4 have their binding sites clustered at ZNF598 promoter and the ZNF598 RP calculated from co-binding sites is significantly negatively correlated with survival in breast cancer patients (Fig. 6e,f). Similar analysis was performed in COAD and we also observed a high association between the target genes of JUND, CEBPB and SRF and the clinical outcomes (Supplementary Fig. 7b). Taken together, these results suggest that the TFs with high CP in a cancer type might function together to cooperatively bind at super-enhancers and form transcriptional condensates to regulate their oncogenic target genes.

## DISCUSSION

The spatial distribution of non-coding regulatory elements in the genome is associated with genome organization and gene regulation, but the spacing patterns of cis-regulatory elements and TF binding sites are rarely studied in a quantitative way. We developed a novel metric, cluster propensity (CP), to survey a large collection of publicly available genomics data, and unraveled the association of the clustered patterns of DNA motif elements and TF binding sites with LLPS transcriptional condensates, which are hypothesized to be the mechanistic basis of super-enhancers^12^. Furthermore, we found that TFs with clustered binding patterns have high liquid-liquid phase separation potentials, directly connecting the genomic pattern to molecular functions. We also found that clustered TF binding sites in cancer cells are highly active and predictive of patient survival. In summary, genomic sequence features and biophysical properties both contribute to the clustered pattern of TF binding, and collectively affect transcriptional condensate formation.

Biomolecular condensates have been a widely studied subject in molecular biology and biophysics. IDR-containing proteins, including many TFs and chromatin regulators, can form large biomolecular condensate through LLPS. In cancer cell nucleus, formation of transcriptional condensates can enhance the genomic targets of oncogenic TFs and induce aberrant 3D chromatin structure for tumor transformation^46, 47^. Principled computational modeling of DNA sequence features has shown that the densely clusters of TF binding sites above sharply defined thresholds can drive the formation of localized condensates to promote enhancer activity and transcription^14^. However, how this sequence pattern occurs in the human genome and how different TFs can induce transcriptional condensates in different cell types are still largely unknown. Our results directly connect genomic information with TFs’ LLPS property, two distinct perspectives that have never been associated before. These results provide quantitative evidence of potential mechanisms of transcriptional condensate formation and super-enhancer activity. In practice, characterization of TF CP and clustered TF binding sites could provide a new approach of studying oncogenic gene regulation and identifying oncogenic drivers in each different cancer type.

We used a data-driven computational approach to reveal the connection between genomic TF binding patterns and LLPS properties. While it provides evidence supporting the hypothesis that transcriptional condensate formation is the mechanism of super-enhancers, we do not have direct experimental data to demonstrate the existence of transcriptional condensate phenomena at super-enhancers, and their dynamic relations with TF binding patterns. Further experiments are needed to validate the formation of transcriptional condensates under the perturbation of identified TFs. Meanwhile, there are other factors missing this work that possibly contribute to the formation of cell type-specific transcriptional condensates, such as long non-coding RNAs, RNA-binding proteins, and genomic DNA and chromatin structure factors that facilitate the chromatin context of condensates. Incorporating these factors in a future updated model will likely improve the characterization of transcriptional condensates’ determinants. Furthermore, in colon cancer and breast cancer case studies, the effects of putative condensate-derived survival predictors are quite different in different cancer types, indicating the complexity of cancer transcriptional regulation and epigenetic mechanisms. Further experiments are required to unravel the cancer type-specific drivers in each individual patient, and to provide translational insights into therapeutic target identification as part of precision medicine practice.

Nevertheless, this work can set a stepstone of future investigations of biomolecular condensates from a genomics perspective.

## METHODS

### Identification of the TF sequence motifs in human genome

DNA sequence motifs in the human genome were searched by FIMO^28^ (v4.12.0) with Jaspar^26^ database (v2018), with a p-value threshold of 1e-4. As a result, 528 TF motifs were included, with a total of 288,687,458 motif sites in the genome, and a median of 551,421 motif sites per motif.

### Public data collection

Super-enhancers (SEs) in 86 samples were collected from the public domain^5^, the chromosomal coordinates were transferred from hg19/GRCh37 to hg38/ GRCh38 using LiftOver^48^. Public ChIP-seq and bigwig profiles were collected from Cistrome Data Browser (DB)^27^. For any TF, only the high-quality peak profiles were used for the subsequent analysis. The quality control thresholds include: FastQC >15, uniquely mapped ratio >0.3, PBC >0.3, FRiP >0.005, 10-fold confident peaks >500, total peaks >2000, and the union DNase I hypersensitive site overlap >0.3, all determined by Cistrome DB.

### Find the nearest site of TFMS/TFBS

The command ‘bedtools closest -D ref -fd -io -t first’ was used to find the distance to the nearest downstream site for each TFMS/TFBS.

### Determination of TFMS CP

For a profile with *N* TF sequence motif matching sites in the human genome, the Poisson point process was used to model the background distribution of the *N* sites randomly occurring in the genome. as 1) the distance of a motif to its downstream motif is independent of the distance of this motif to its upstream motif, 2) the average distance between two motifs is *L*/(*N*+1), where *L* is the total length of the human genome, 3) the two motifs cannot occur at the same location. The TFMS CP is derived from the statistic of two-sided Kolmogorov-Smirnov (K-S) test by comparing distribution of log10 distances to the down-stream motif for a TF sequence motif profile (T) and genomic background control (C) as follows:

1, A is defined as the statistic of K-S test following the null hypothesis that Log_10_Distance (T) < Log_10_Distance (C).
2, B is defined as the statistic of K-S test following the null hypothesis that Log_10_Distance (T) > Log_10_Distance (C)..
3, CP is determined as 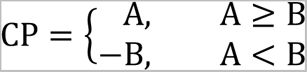

### Fitting of TFMS with Gamma distribution

For each TF motif profile, the Gamma(*k*, *θ*) distribution, where *k* is shape parameter and *θ* is the scale parameter were used to fit the distribution of TFMS in the genome. *θ* is determined as the genome length divided by the number of motifs. The estimated k from all TFs were displayed in Supplementary Fig. 1d.

### Determination of TFBS CP

For a TF ChIP-seq profile with *N* peaks, the same number of *N* motif sites for the same factor were randomly selected in the genome as the background control. As described in the **Determination of TFMS CP** section, a CP is derived from the two-sided K-S test by comparing the distribution of log10 distances to the down-stream site from a TF ChIP-seq binding profile (T) and the control (C). The random selection of the background control was performed 100 times and the average of 100 CPs was use for the TFBS CP of the ChIP-seq profile, i.e., the TFBS CP of the factor in the corresponding cell type. For a factor with multiple ChIP-seq profiles from the same cell type, the average of TFBS CPs across all ChIP-seq profiles was used as the TFBS CP of the factor in the cell type. To get the normalized cell-type-specific CP of a factor in a cell type, the TFBS CP scores of the factor in all cell types were collected for z-score normalization, and the normalized TFBS CP of the factor in the corresponding cell type was shown in the x-axis of Fig. 2e. For each cell type, the TFs were ranked by the average rank of CP and z-score normalized CP. The top5 TFs were highlighted in Fig. 2e, and the rankings were displayed in Fig. 4b-e.

### Enrichment of TFMS at union SEs

For each TFMS profile, the two-tailed Fisher’s exact test was applied to test the enrichment of TFMS at the union of SEs from 86 samples using the randomly selected genomic loci as control. Odds ratio (OR) >1 (log2 OR >0) indicating the TFMS are more enriched at union SEs compared to the genomic background control (Fig. 1d). P-values were calculated using the Fisher’s exact test.

### Identification of clustered and non-clustered TFBS

To identify the clustered- and non-clustered (C-/NC-) TFBS from a TF ChIP-seq profile, the genomic background control is first selected as randomly selected the same number of sequence motifs from the same factor. The distribution of distances to the down-stream sequence motif were collected from the control and the 5-th percentile distance/score was kept. All the 5-th percentile scores from 100 random samples of background control were averaged as the cutoff for C-TFBS and NC-TFBS. TFBS with a neighbor less than the cutoff were grouped into C-TFBS as the binding sites are significantly close to each other compared to the randomly selected control, while other TFBS were groups into NC-TFBS as those sites do not have significantly closed neighbors. C-TFBS for each TF ChIP-seq profile were merged as “bedtools merge -d 5-th-cutoff”. For TFs with multiple ChIP-seq profiles in a same cell type, the C-TFBSs were further merged across all ChIP-seq profiles as the C-TFBSs of the TF in the cell type, and all NC-TFBS excluding C-TFBS were merged across all ChIP-seq profiles as NC-TFBS.

### Enrichment of C-TFBS at cell-type-specific SEs

For each TF and each cell type, the two-tailed Fisher’s exact test was applied for the enrichment of C-TFBS at the cell-type-specific SEs using the NC-TFBS as control. Odds ratio (OR) >1 (log2 OR >0) indicating the C-TFBSs are more enriched at cell type-specific SEs compared to NC-TFBS (Fig. 4b,c, Supplementary Fig. 5a).

### ATAC-seq regulatory potential on TFBS

We use the TCGA ATAC-seq bigwig profiles from primary patients^41^ to calculate the chromatin accessibility regulatory potential (RP)^42^ at TFBSs (Fig. 4a). For each TFBS, the chromatin accessibility RP was calculated as the sum of ATAC -seq levels weighted by the genomic distance from the peak center. Specifically, ATAC-seq levels surrounding peak *i* were collected and weighted by an exponential decay function for the total chromatin accessibility *RP_i_* on this peak:

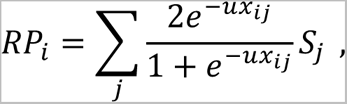

Where *S_j_*_’_ is the chromatin accessibility level surrounding peak *i* (peak center +/-100kb), and *x_ij_*_’_ is the distance between the center of peak *i* and *S_j_*. The parameter *u* determines the decay rate and is set so that the half-life of the decay function is 10kb. The ATAC-seq RPs comparing C-TFBSs and NC-TFBSs were assessed using two-sided t-test and the statistics and p-values were shown in Fig. 4b,c, Supplementary Fig. 5a.

### Differential ATAC-seq analysis

We used the processed data from Ref.^41^ that include a matrix of normalized ATAC-seq insertion counts within the TCGA pan-cancer peak set to assess the differential chromatin accessibility at each ATAC-seq peak. The differential ATAC-seq score at each peak was defined as the two-sided t-test statistics comparing ATAC-seq levels from patients in the corresponding cancer type vs. patients from other cancers (Fig. 4a). The differential ATAC-seq scores comparing C-TFBSs and NC-TFBSs were assessed using two-sided t-test and the statistics and p-values were shown in Fig. 4b,c, Supplementary Fig. 5a.

### Chromatin interactions

Hi-C data were processed using HiC-Pro^49^. Contact maps were generated at a resolution of 5kb and BART3D^45^ was applied on the raw count matrices for normalization. The chromatin interactions with surrounding genomic loci (<100 kb) were collected at each TFBS. The interactions scores comparing C-TFBSs and NC-TFBSs were assessed using two-sided t-test and the statistics and p-values were shown in Fig. 4b,c, Supplementary Fig. 5a.

### Identification of differential chromatin interactions

Hi-C data were first processed using HiC-Pro^49^. Contact maps were generated at a resolution of 5kb. BART3D^45^ was applied on raw count matrices between samples before and after RAD21 degradation in HCT-116 cells to generate genome wide differential chromatin interaction (DCI) profiles (--genomicDistance 100000). DCI score at each 5kb bin was then mapped to the TFBS to infer the differential chromatin interactions at the binding site (Fig. 4b,c, Supplementary Fig. 5a).

### Detection of mutation at TFBS and genes encoding the TFs

We use the whole genome sequencing (WGS) data from the International Cancer Genome Consortium (ICGC)^44^ to check the mutations at TFBS and genes that encoding the TFs. For each TFBS in a cell type, the mutation rate at the sequence motif within the TFBS was calculated as the occurrence of mutation events across all patient samples from the matched cancer type divided by the total patient numbers. The mutation rates for C-TFBS and NC-TFBS were then averaged over the number of binding sites and shown in Fig. 4d,e, Supplementary Fig. 5b.

For each TF, the mutation rate at the gene that encoding the TF were assessed the same way as the TFBS. The patient samples were separated into two groups by the ATAC-seq RPs at C-TFBS from the corresponding TF for each cancer type, and the mutation rate of the genes encoding the TF were compared between patients with higher RP and lower RP and were shown in Supplementary Fig. 6a.

### Determination of TFBS target genes

For a set of TFBSs, either selected as the C-TFBS from a TF or the co-binding sites shown in Fig. 6a, the ATAC-seq peaks that overlapped with the TFBSs were used to calculate the regulatory potential (RP)^42^ on each gene. The ATAC-seq peak levels surrounding gene *i* (TSS +/-100kb) were collected and weighted by an exponential decay function as shown above, e.g., for the *RP_i_* on gene *i*, *S_j_* is the ATAC-seq peak level and *x_ij_* is the distance between TSS of gene *i* and ATAC-seq peak *j*. The parameter *u* determines the decay rate and is set so that the half-life of the decay function is 10kb (Fig. 6c).

### Survival analysis

Univariate survival analysis at each ATAC-seq peak in each cancer type was applied using patient samples with both supported TCGA clinical data and ATAC-seq profiles^41, 50^. For each selected cancer type and each identified ATAC-seq peak, the primary patients were separated into two equal-sized groups based on the chromatin accessibility at the ATAC-seq peaks (top 50% and bottom 50%). The Kaplan-Meier (K-M) method was used to create the survival plots and log-rank test was used to compare the differences of survival curves.

Univariate survival analysis at each gene for each cancer type was applied using patient samples with TCGA clinical data and ATAC-seq profiles. For each selected cancer type and each gene, the patient samples were separated into two equal-sized groups based on the RP calculated from TFBS overlapped ATAC-seq peaks. The K-M method was used for the survival plots and log-rank test was used to compare the differences of survival curves for the p-values.

## DATA AND CODE AVAILABILITY

Re-analyzed data results, software packages developed for Cluster Propensity calculation, and all codes and scripts to produce the results are available at: https://github.com/zang-lab/transcriptional_condensates

## ACKNOWLEDGEMENTS

The authors thank Dr. Hao Jiang for helpful discussions. This work was supported by US National Institutes of Health grant R35GM133712 to C.Z.

---

**Supplementary Fig 1.**
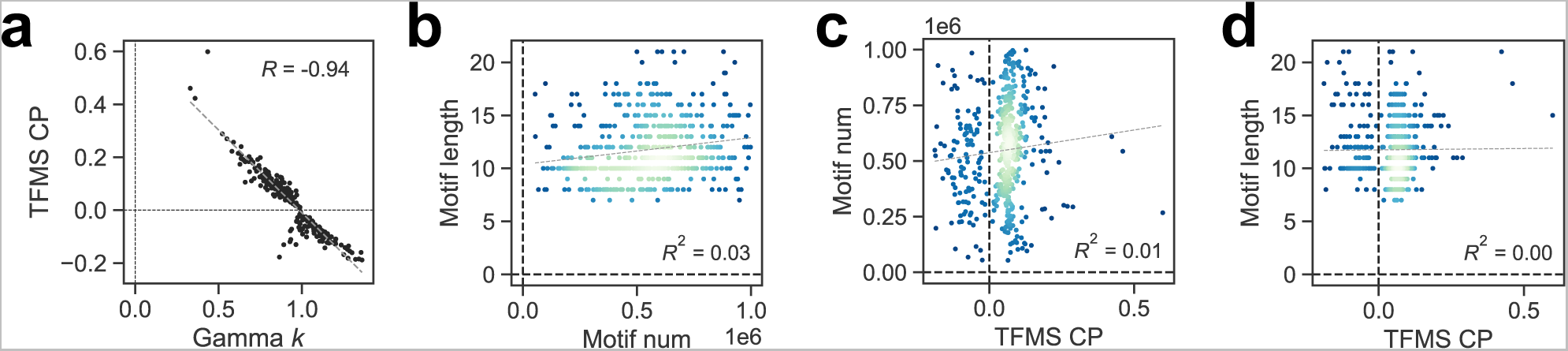
Different TFs show different TFMS CPs. **(a)** Association of Gamma k with TFMS CP. **(b-d)** Scatter plots of correlation among TFBS CP, number and length of TF motifs.

**Supplementary Fig 2.**
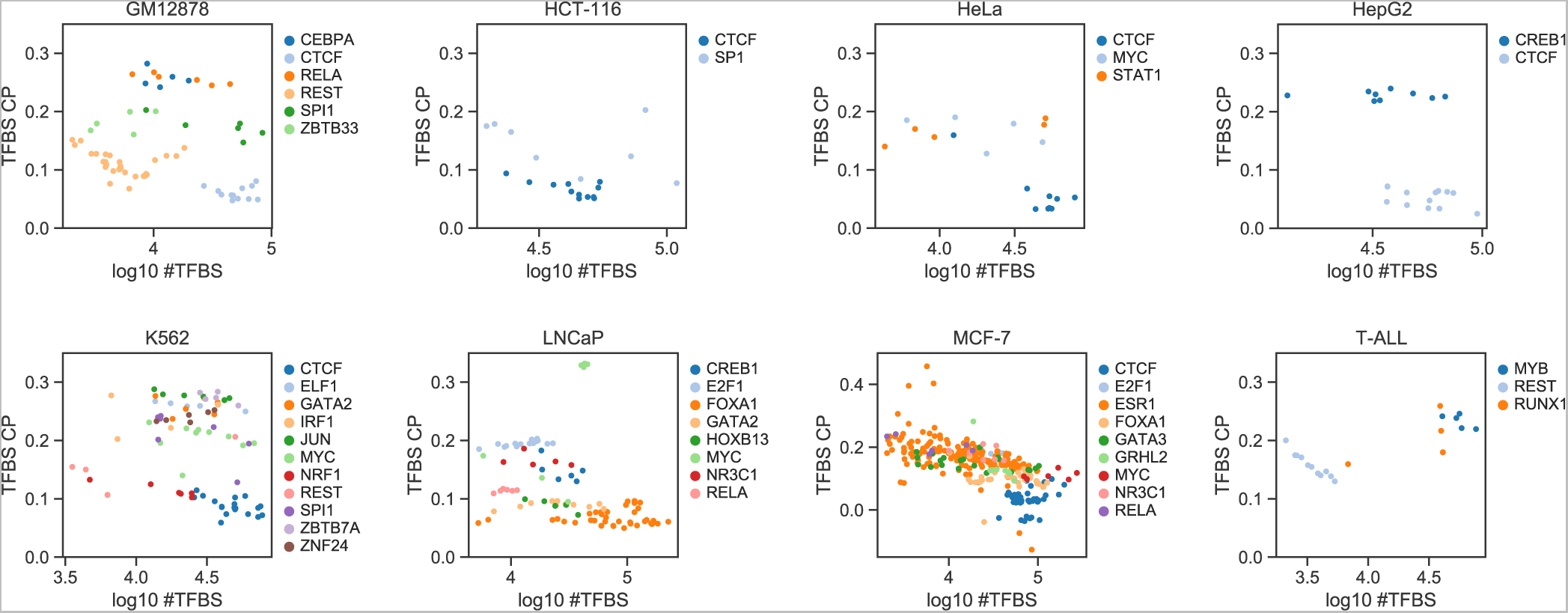
TFBS CPs are not correlated with the number of peaks in the ChIP-seq profiles. Scatter plots of TFBS CP (y-axis) against the number of binding sites (log10) in ChIP-seq profile in each of the 8 cell types with at least 5 TFs having ChIP-seq data.

**Supplementary Fig 3.**
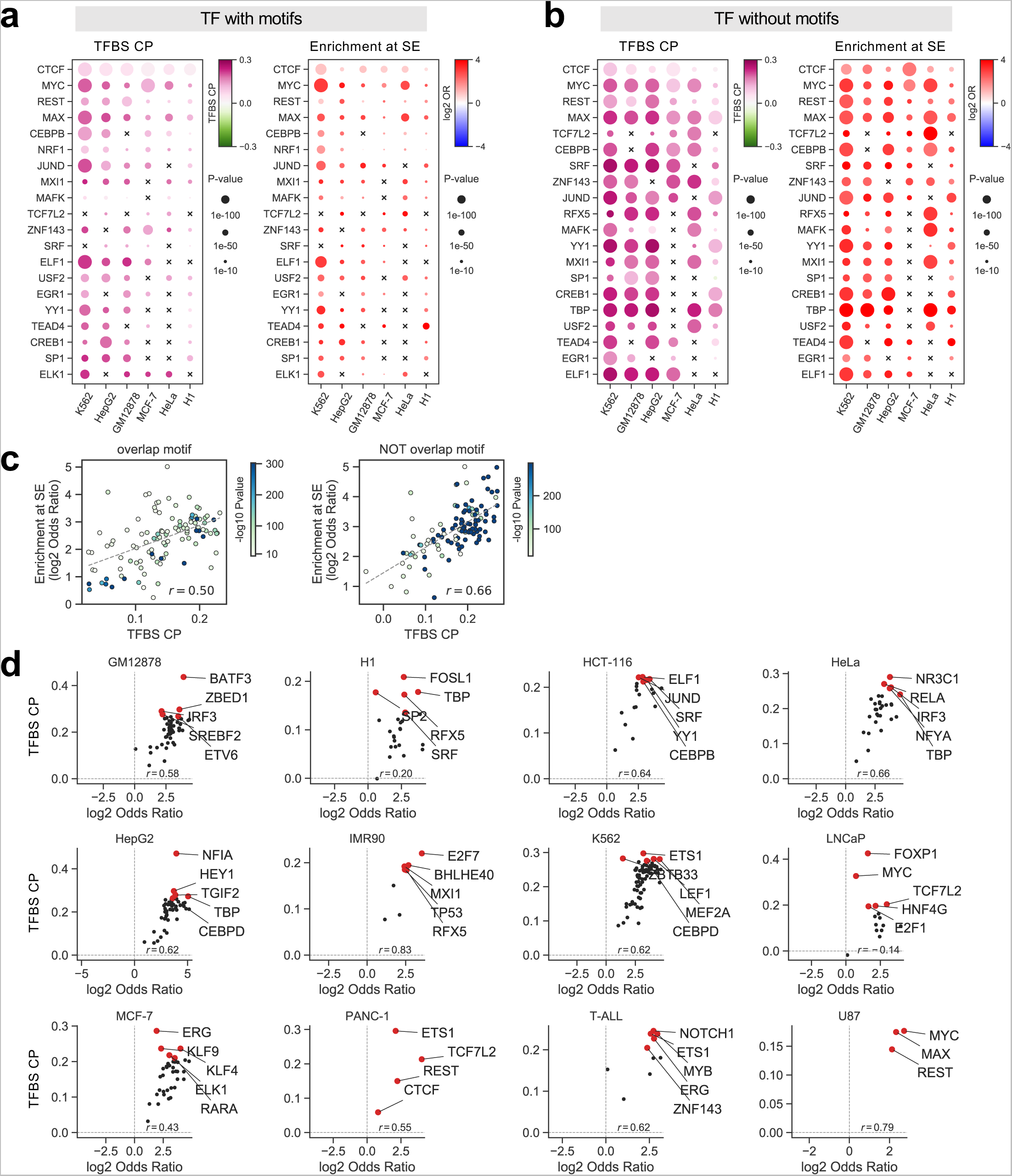
TFs show different TFBS CPs in different cell types. **(a,b)** TFBS CP (left) and enrichment of TFBS at cell type-specific SE compared with random control (right) of 20 TFs in 6 cell types for TFBS with motif (a) and without motif(b). **(c)** Scatter plots of correlation of TFBS CP (x-axis) and enrichment of TFBS at cell type-specific SE compared with random control (y-axis) of 20 TFs in 6 cell types. **(d)** Scatter plots of TFBS CP (y-axis) against the enrichment of TFBS at cell type-specific SE (x-axis) in each of the 12 cell types with at least 3 TFs having ChIP-seq data.

**Supplementary Fig 4.**
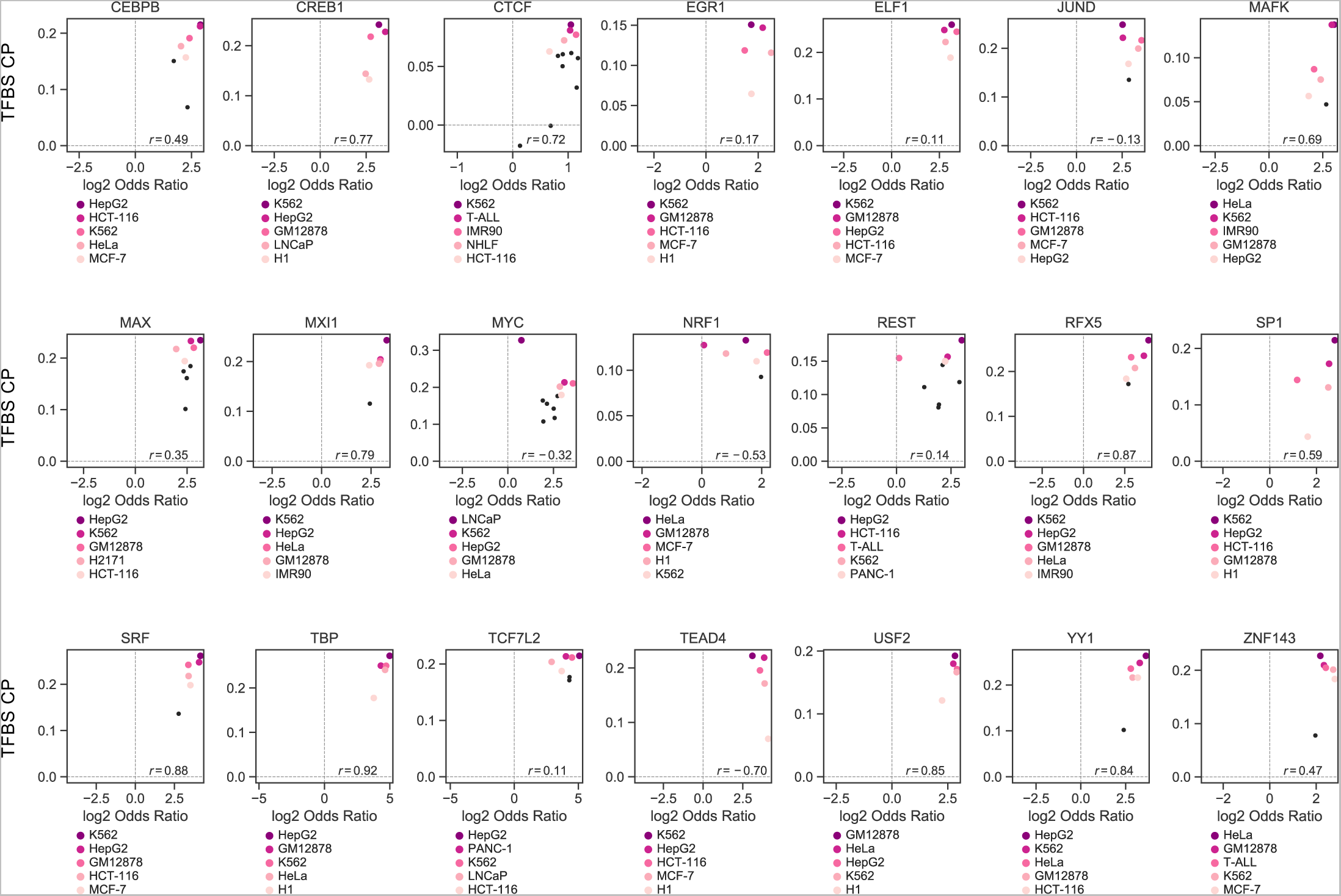
The same factor has different TFBS CPs across different cell types. Scatter plots of TFBS CP (y-axis) against the enrichment of TFBS at cell type-specific SE (x-axis) in each of the 21 factors with at least 5 cell types having ChIP-seq data.

**Supplementary Fig 5.**
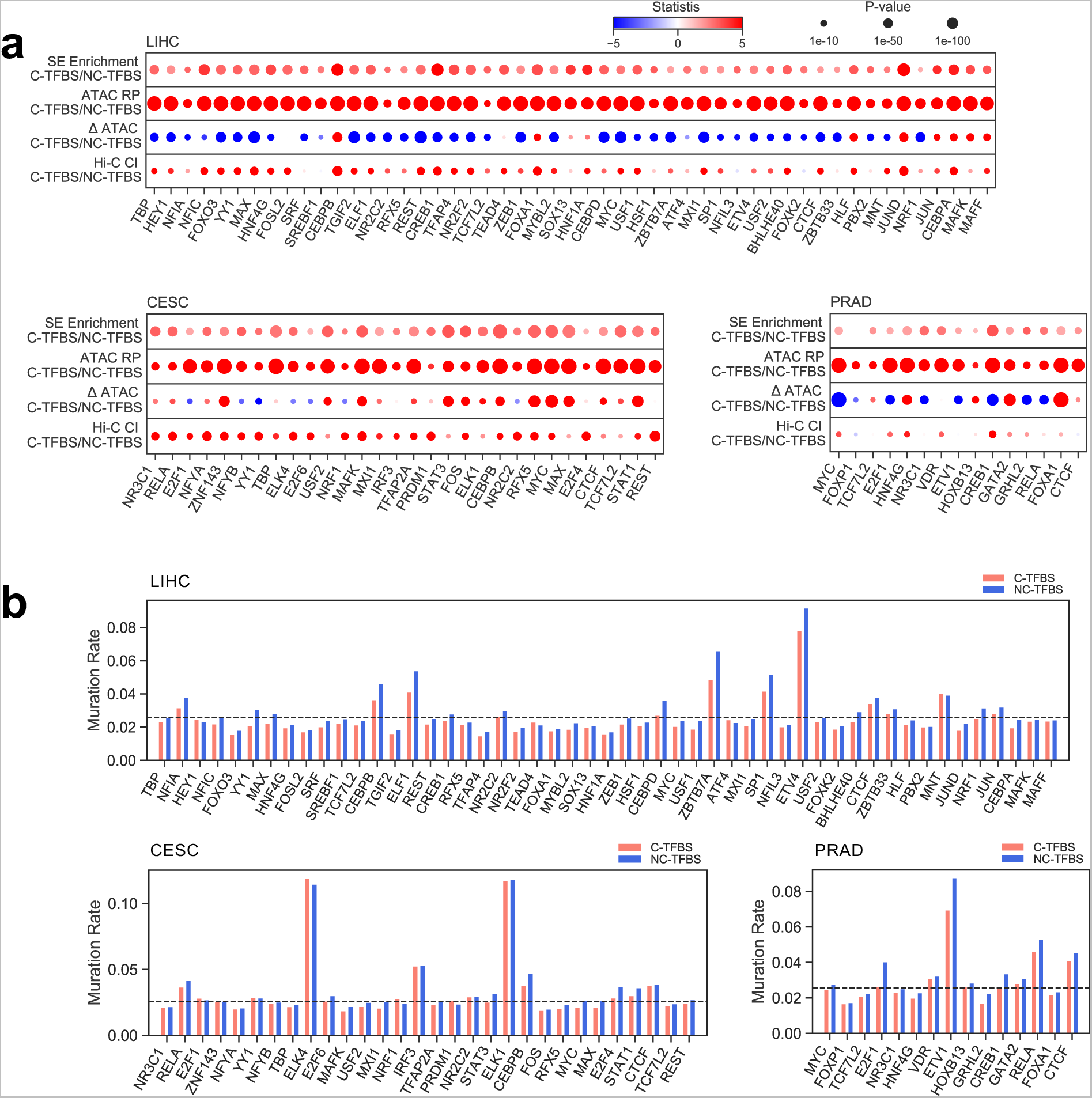
Chromatin activity and mutations of C-TFBS and NC-TFBS in different cancer cells. **(a)** The comparison of enrichment at cell-type-specific SEs, ATAC-seq RP, differential ATAC-seq score and Hi-C chromatin interactions between C-TFBS and NC-TFBS in LIHC, CESC and PRAD. TFs were ranked on x-axis by CP rank as shown in Fig. 2e. **(b)** Mutation rate at motif loci within the binding sites comparing C-TFBS and NC-TFBS in LIHC, CESC and PRAD. TFs were ranked on x-axis by CP rank as shown in Fig. 2e.

**Supplementary Fig 6.**
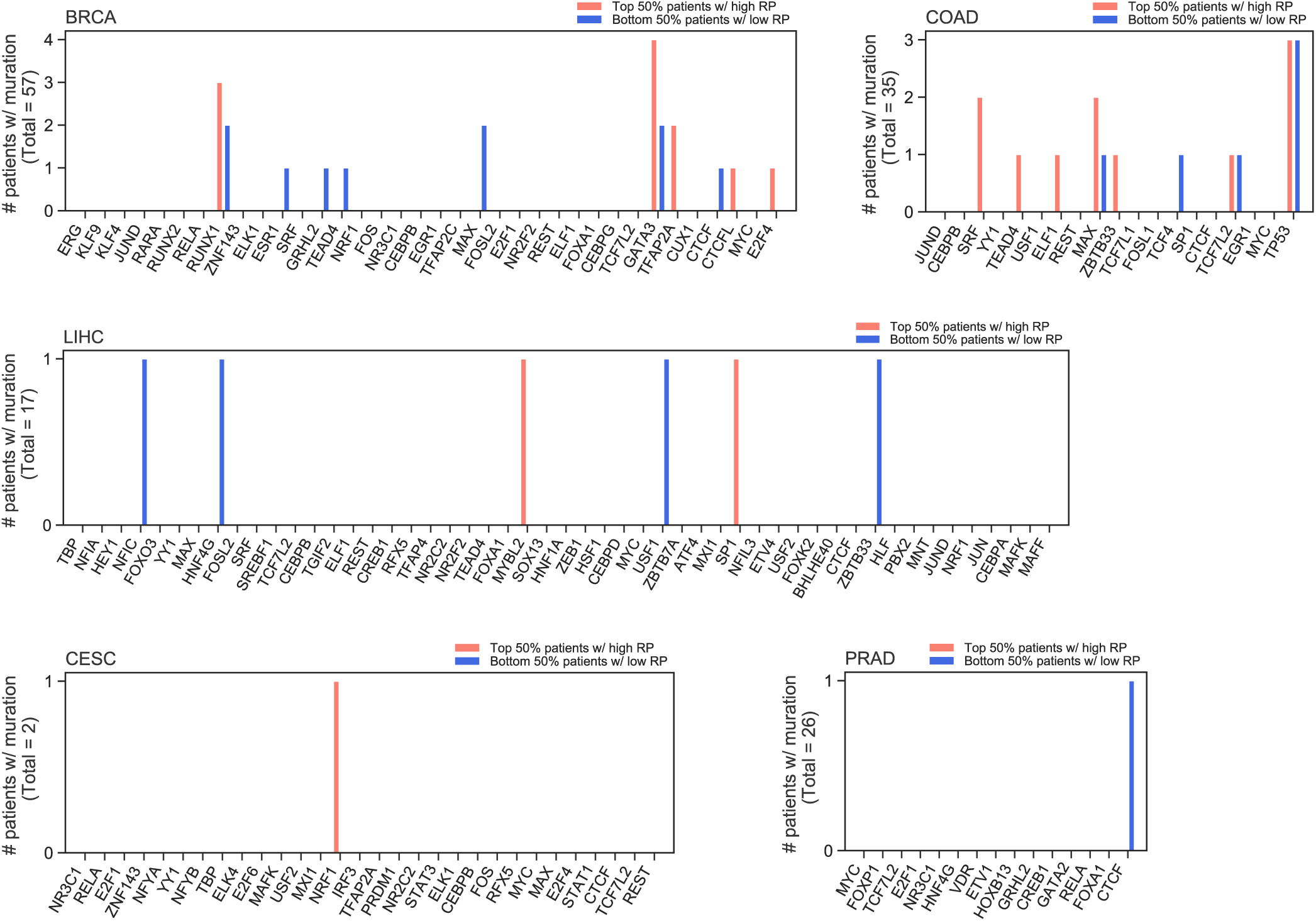
Mutations at genes encoding TFs in different cancer cells. The mutation rate of genes encoding the TFs in LIHC, CESC and PRAD. For each factor and in each cell type, the patients were evenly separated into two groups by their averaged ATAC-seq RP at the C-TFBSs from the corresponding TF. TFs were ranked on x-axis by CP rank as shown in Fig. 2e.

**Supplementary Fig 7.**
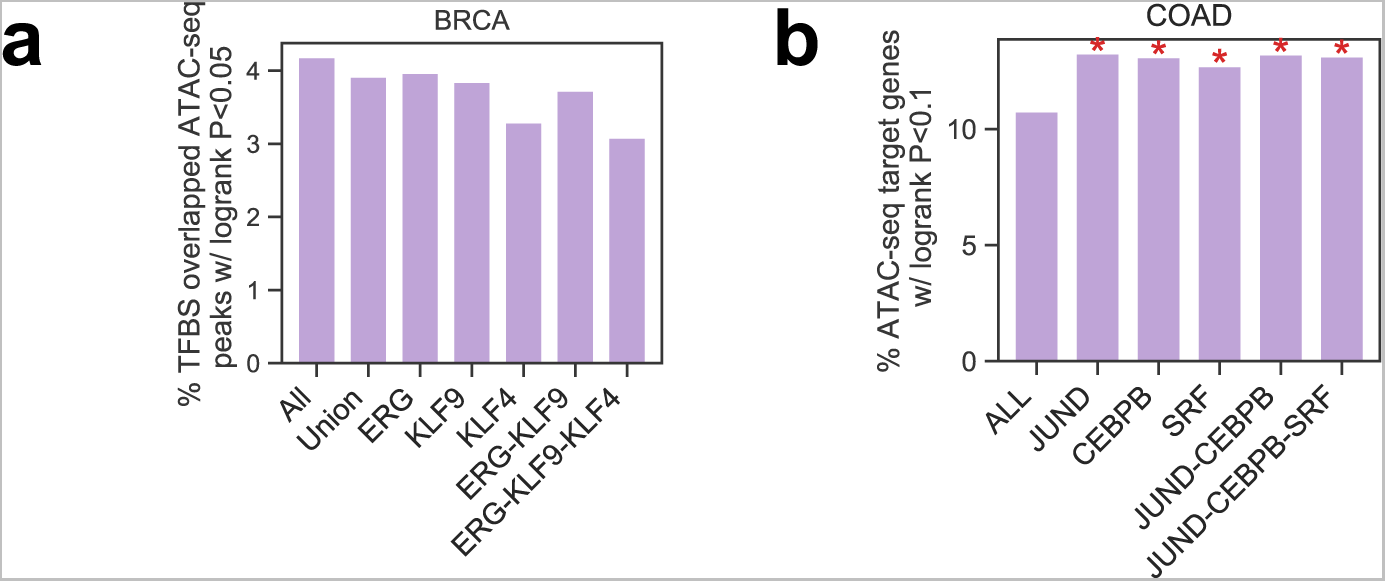
Association of chromatin accessibility levels at clustered TFBSs and clinical outcome. **(a)** Bar plot of percentage of clinical associated ATAC-seq peaks overlapping binding and co-binding of C-TFBS of the 3 factors with the highest CP rank in BRCA. **(b)** Bar plot of percentage of clinical associated target genes of the 3 factors with the highest CP rank in in COAD.

## Notes

### Competing Interest Statement

The authors have declared no competing interest.

